# Infrapatellar Fat Pad Extracellular Vesicles Induce a Pro-Angiogenic VEGFA^high^/BMP4^low^ Switch in Articular Chondrocytes: Implications for Chondrosarcoma

**DOI:** 10.64898/2026.08.25.746948

**Authors:** Joshua M. J. Price, Caitlin Ditchfield, Hussein Farah, Bethy Airstone, Natalie Lachlan-Jiraskova, Edward T. Davis, Simon W. Jones

## Abstract

Chondrosarcoma is a hyper-vascularised, chemoresistant cartilage malignancy driven by VEGF-centred angiogenesis, and local adipose depots are increasingly recognised as paracrine drivers of tumour angiogenesis via adipokines and extracellular vesicles (EVs). The infrapatellar fat pad (IFP), an inflammatory adipose depot within the articular joint in direct cartilage contact, is a key local source of adipose-derived EVs, and thus a candidate driver of angiogenesis in chondrosarcoma. The aim of this study was to determine whether the IFP is a productive source of EVs, and whether IFP-derived EVs induce angiogenesis in articular chondrocytes. The IFP released significantly more EVs than subcutaneous fat (n = 8 per depot; p = 0.027). Treating primary human articular chondrocytes with IFP EVs for 24 h upregulated VEGFA (+1.6-fold, p = 0.036) and downregulated BMP4 (−2.4-fold, p = 0.011), engaging the VEGF/eNOS/ERK axis that drives chondrosarcoma angiogenesis. Re-analysis of a previously published phospho-kinase dataset from the same donor EVs, corroborated by a pooled donor-group analysis (n = 3), supported activation of eNOS, ERK1/2, PLC-γ1 and HSP27. These findings identify the IFP as a dominant source of EVs within the articular joint, which can induce a pro-angiogenic, VEGF-axis switch in articular cartilage cells, supporting a signalling model relevant to chondrosarcoma angiogenesis.

## 1. Introduction

Chondrosarcoma is the second most common primary bone malignancy and is characteristically resistant to chemotherapy and radiotherapy, leaving surgical resection as the primary treatment option [1]; its progression is closely tied to pathological angiogenesis, since the cartilage from which it arises is normally avascular and neo-vascularisation is required to support tumour growth and invasion [2]. VEGF-A-dependent angiogenesis is a well-established driver of this process [2]: adipokines such as adiponectin and resistin have each been shown to promote VEGF-A-dependent angiogenesis in human chondrosarcoma cells via PI3K/Akt/mTOR/HIF-α and related signalling [3,4]. This positions the local adipose tissue surrounding cartilage and bone as a plausible paracrine source of angiogenic signal in the chondrosarcoma microenvironment. However, how local joint adipose depots communicate with articular cartilage cells to drive this angiogenic switch is not well defined.

Adipose depots communicate with neighbouring and distant tissues largely through their secretome, a mixture of soluble adipokines and extracellular vesicles (EVs) that carry protein and RNA cargo between cells [5]. Beyond its soluble adipokine component, this secretome, and its EV fraction in particular, can drive pro-angiogenic paracrine effects in other tissue contexts [6], and EV-mediated crosstalk between adipose tissue and tumour cells has been proposed as a driver of angiogenesis and progression in other cancers, positioning EVs as a candidate active constituent of the wider adipose secretome. The infrapatellar fat pad (IFP) is a candidate local source of this signal: unlike subcutaneous fat, it sits inside the joint capsule in direct contact with synovium and cartilage, and has a demonstrably inflammatory phenotype relative to subcutaneous fat [7], including elevated secretion of IL-6 and its soluble receptor [8]. The same angiogenic, VEGF-centred switch implicated in chondrosarcoma is also a feature of degenerative cartilage disease, where pathological neo-vascularisation brings inflammatory cells and matrix-degrading proteases into cartilage that is normally avascular, and gives sensory nerves a scaffold to grow into cartilage that is not normally innervated, contributing to structural damage and chronic joint pain [9]. However, whether the IFP is a productive source of EVs, and whether IFP-derived EVs are capable of modulating the expression of angiogenic mediators in human articular chondrocytes has not been established.

## 2. Methods

### 2.1 Ethics and donors

Adipose tissue and articular cartilage were collected peri-operatively from adults undergoing elective orthopaedic surgery for osteoarthritis at the Royal Orthopaedic Hospital (Birmingham, UK) or Russell’s Hall Hospital (Dudley, UK), with written informed consent, under UK National Research Ethics Service approval 16/SS/0172. Joint IFP was the primary adipose tissue studied; paired subcutaneous fat was collected for the depot comparison (Section 3.1).

### 2.2 Conditioned media, EV isolation and particle characterisation

Adipose tissue was minced (∼2–3 mm) and incubated in DMEM (1 g : 10 mL) for 24 h (37°C, 21% O_2_, 5% CO_2_) to generate adipose-conditioned media (ACM), clarified (2,000 × g, 20 min, ×2) and 0.22 µm-filtered. Chondrosarcoma-conditioned media (CCM) was generated identically from serum-starved SW1353 cells (ATCC).

Polymer-based precipitation was performed on raw conditioned media using the EV Precipitation Solution (Cell Culture; Novus Biologicals). Raw conditioned media was pre-cleared by sequential centrifugation (300 × g, 10 min; 1,200 × g, 20 min; 10,000 × g, 30 min), mixed 1:1 with EV Precipitation Solution, incubated on ice for 1 h, then centrifuged (10,000 × g, 20 min) to pellet EVs. A final 1,500 × g, 2 min spin was done to remove residual supernatant. The pellet was resuspended in 100 µL PBS for NTA, alongside a matched raw-material (no-precipitation) control.

The 24 h-vs-48 h ACM collection-time optimisation and the ultracentrifugation (UC) isolation validation against raw media using SW1353 CCM were performed by nanoparticle tracking analysis (NTA; NanoSight Pro, Malvern Panalytical).

EVs used in all subsequent experiments were isolated by ultracentrifugation (100,000 × g, 16 h, 4°C). The fat pad-versus-subcutaneous EV yield comparison (Section 3.1) was performed independently by nanoscale flow cytometry (CytoFLEX nano, Beckman Coulter) gated on violet side scatter with fluorosphere calibration and swarm-detection controls, following a dilution-linearity validation (R^2^ = 0.997), consistent with established performance characteristics of the platform [10], MISEV2023 reporting guidance [11], and the MIFlowCyt-EV framework for standardised reporting of extracellular vesicle flow cytometry experiments [12] (Table S1, Supplementary Materials).

### 2.3 Cell treatment and RT^2^ Profiler array

Primary human chondrocytes were isolated from articular cartilage by collagenase digestion (Sigma C9891, 2 mg/mL, 5–15 h, 37°C), filtered (70 µm) and cultured in chondrocyte growth medium. Chondrocytes were used for experiments before passage 2 to retain their phenotype. Chondrocytes were treated for 24 h with the UC-isolated fat pad EV fraction (resuspended to the concentration present in ACM; or left untreated), and SW1353 cells were profiled in parallel.

RNA was extracted following the manufacturer’s protocol (RNeasy, Qiagen) and profiled by RT^2^ Profiler Human Angiogenesis & Inflammatory Cytokines array (PAHS-024Z, 84 targets; QuantStudio 6 Flex). Per-sample Cт values were normalised to the mean of five reference genes (ACTB, B2M, GAPDH, HPRT1, RPLP0); fold-changes and two-sample t-tests were computed directly from the ΔCt/2^-ΔCt values (n = 3 per group). Principal component analysis and hierarchical clustering were performed on z-scored 2^-ΔCt expression across all 84 targets.

### 2.4 Phospho-kinase array

Phospho-kinase array data were reanalysed from our previously published dataset [18], generated using human OA primary cells (myotubes differentiated from skeletal-muscle-derived myoblasts of patients from the same elective-orthopaedic-surgery recruitment pathway as this study) treated for 24 h with fat pad EV fractions from normal-weight, overweight or obese donors (one pooled sample per BMI group), or left untreated. Cells were lysed on ice with RIPA buffer supplemented with protease and phosphatase inhibitors (cOmplete and PhosSTOP tablets, Roche) and screened on a 16-target Proteome Profiler phospho-kinase array (ARY003C, R&D Systems), as described previously [18]. Here we reinterpret this dataset through the lens of VEGF-associated signalling, using the KEGG VEGF pathway mapping established in Section 3.2.

### 2.5 Statistics

Two-group comparisons used Student’s/Welch’s t-test for normally distributed data (Shapiro-Wilk, p > 0.05) or the Mann-Whitney U test otherwise. The IFP-versus-subcutaneous EV comparison (Section 3.1) used a one-sided test, since IFP EV yield was predicted a priori to exceed subcutaneous, given its established inflammatory phenotype [7,8]. RT^2^ Profiler array p-values are nominal and uncorrected, reflecting its use as a hypothesis-generating screen. Significance was set at p < 0.05.

## 3. Results

### 3.1 The infrapatellar fat pad is a dominant adipose EV source

By nanoscale flow cytometry, EV concentration was significantly higher in IFP ACM than in subcutaneous ACM from the same donors (n = 8 per depot; mean 1.42 × 10^11^ vs 5.66 × 10^10^ particles/mL; one-sided Student’s t-test p = 0.027) (Figure 1A). Combined with its established pro-inflammatory phenotype, this makes the IFP both the more inflammatory and the more EV-productive depot adjacent to cartilage, and a relevant adipose EV source for this comparison with cartilage-adjacent tissue. Alongside this depot comparison we ran the method-optimisation experiments that underpin it, using nanoparticle tracking analysis (NTA). ACM collection time (24 h vs 48 h) did not alter EV concentration or size (Figure 1B, C; n = 3, p = 1 for both), supporting the 24 h collection window used throughout. Ultracentrifugation (UC) isolation was also validated against raw conditioned media using SW1353 chondrosarcoma-conditioned media: UC gave a significantly higher particle concentration (Figure 1D; n = 5 repeated NTA captures of one raw preparation and its corresponding UC pellet; Mann-Whitney U p = 0.016) without significantly altering particle size (Figure 1E; p = 0.222). We also trialled a polymer-based precipitation kit (Novus Biologicals) as a candidate alternative isolation method: precipitation did not significantly change particle concentration relative to raw material (Figure 1F; Wilcoxon p = 0.29) but did significantly increase particle size (Figure 1G; Wilcoxon p = 0.0029), so precipitation was not carried forward as an isolation method for this study.

**Figure 1.**
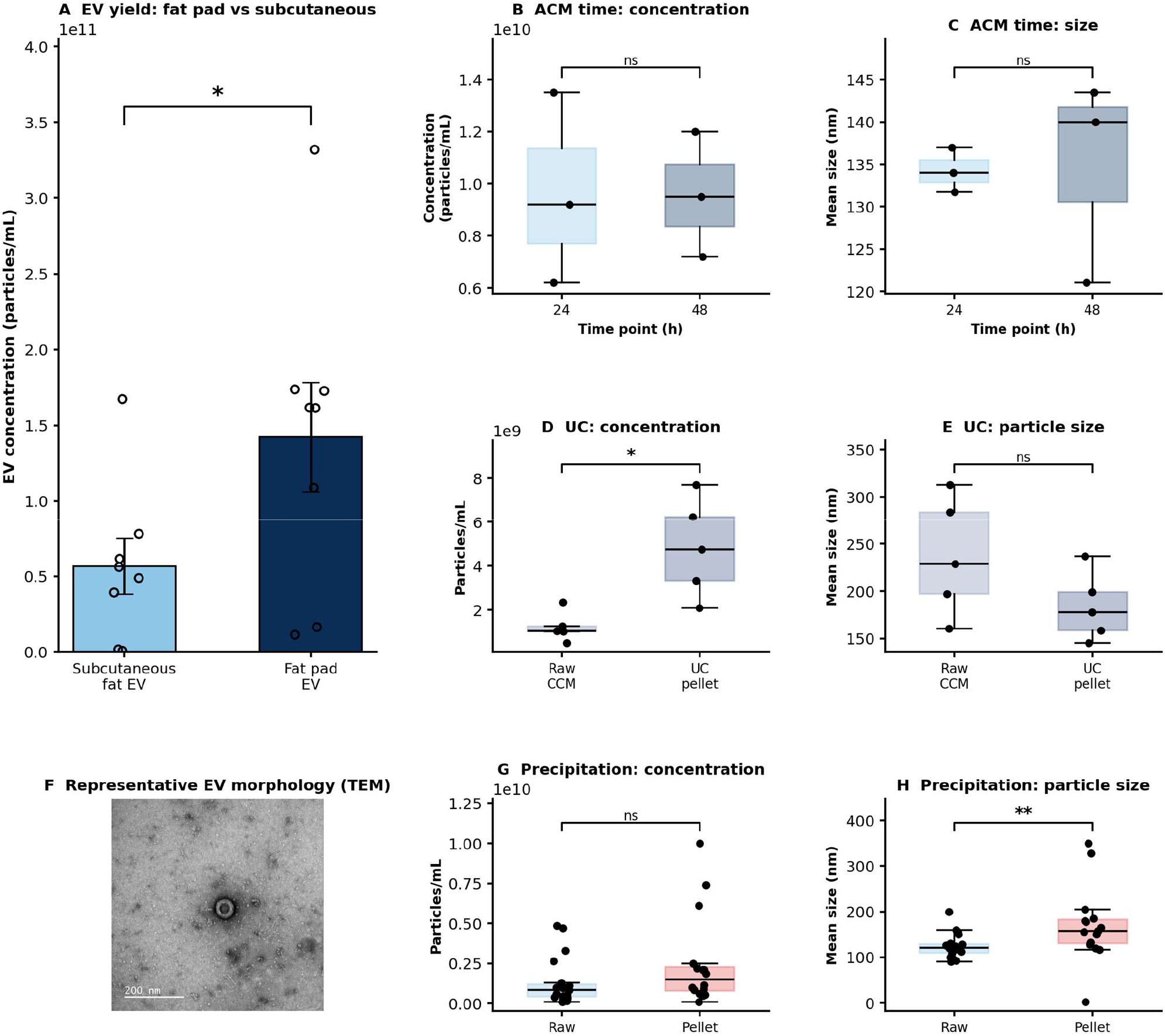
The IFP releases more EVs than subcutaneous fat, method optimisation, and EV morphology. (A) EV concentration by nanoscale flow cytometry (CytoFLEX nano) in ACM from subcutaneous fat versus IFP; n = 8 per depot; one-sided Student’s t-test p = 0.027. (B, C) ACM collection time: EV concentration and mean particle size by nanoparticle tracking analysis (NTA; NanoSight Pro, Malvern Panalytical) after 24 h vs 48 h of tissue conditioning (n = 3; p = 1 for both), supporting a 24 h collection window. (D, E) Ultracentrifugation (UC) validation by NTA using SW1353 chondrosarcoma-conditioned media (CCM): a single raw CCM preparation and its corresponding UC pellet were each measured across 5 repeated NTA captures (technical replicates, not independent biological samples). UC pellet concentration was significantly higher than raw CCM (D; Mann-Whitney U p = 0.016) without a significant change in particle size (E; p = 0.222), supporting UC as the isolation method used throughout. (F) Representative transmission electron microscopy (TEM) image of a UC-isolated fat pad EV, negatively stained with uranyl acetate, showing a spherical structure enclosed by a lipid bilayer membrane, consistent with the expected morphology of EVs (imaging protocol and description as reported previously [18]; scale bar, 200 nm). (G, H) Polymer-based precipitation (Novus Biologicals) trialled as a candidate alternative isolation method, raw material versus precipitated pellet: precipitation did not significantly change particle concentration (G; Wilcoxon p = 0.29) but significantly increased particle size (H; Wilcoxon p = 0.0029), so this method was not adopted. Box, IQR; whiskers, range; points, individual replicates/captures.

### 3.2 IFP EVs upregulate chondrocyte VEGFA within a focused, non-tumour-like response

We profiled untreated chondrocytes, IFP EV-treated chondrocytes and the SW1353 chondrosarcoma line on the 84-gene RT^2^ Profiler array (Figure 2). Principal component analysis and hierarchical clustering placed SW1353 in a distinct transcriptional state, separated from primary chondrocytes along the first principal component (40% of variance), while IFP EV-treated chondrocytes clustered closer to untreated chondrocytes (Figure 2A, B). SW1353 was characterised by higher immune/cytokine genes (CD70, FAM3B, SPP1, IL15, IL21) and markedly lower cartilage/skeletal differentiation genes (GDF5, TNFRSF11B, BMP2, INHBA, BMP4) than chondrocytes (Figure 2C, E), confirming that the tumour line occupies a different transcriptional space from healthy chondrocytes.

**Figure 2.**
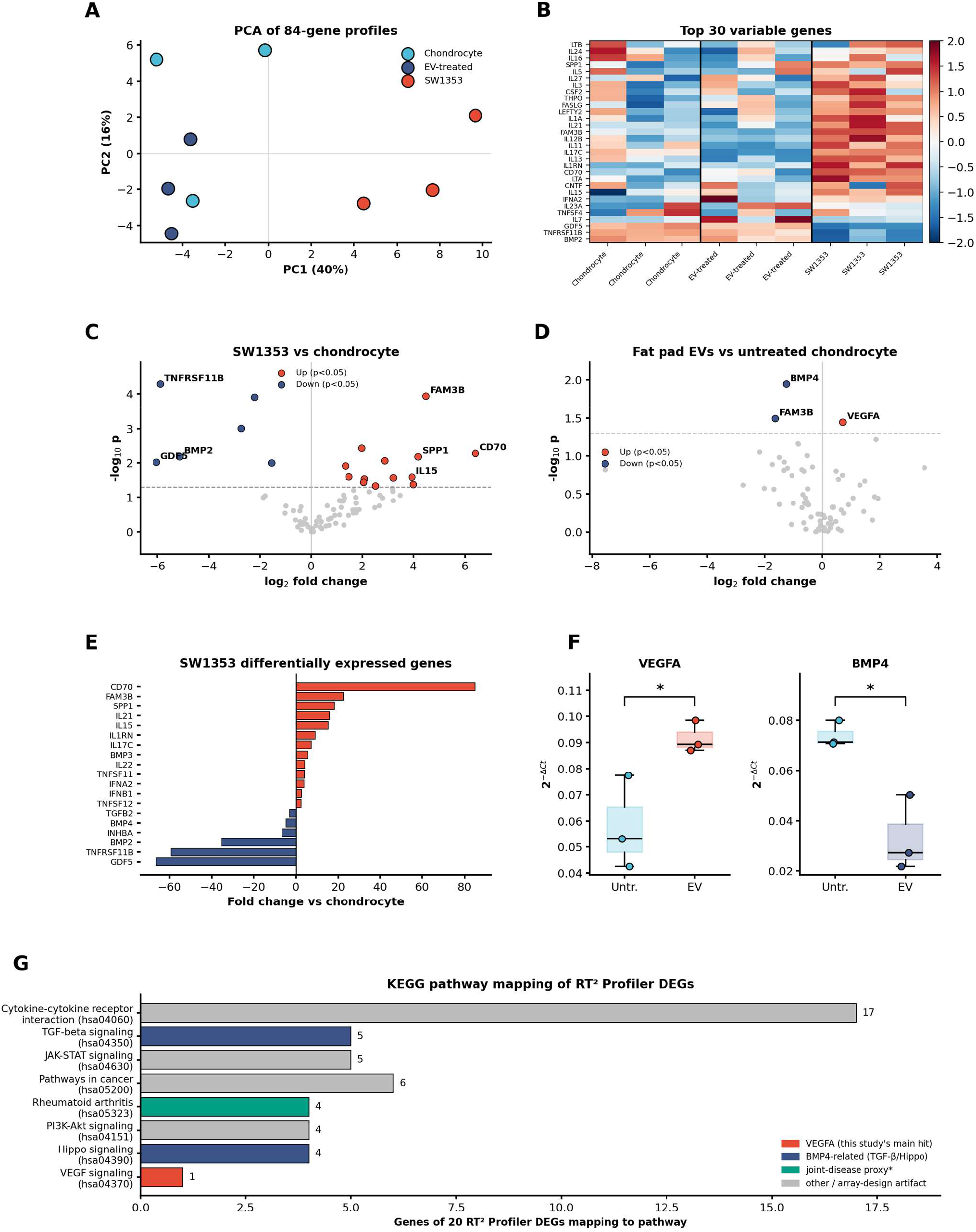
Transcriptional response of chondrocytes to IFP EVs, RT^2^ Profiler array, and KEGG pathway mapping. (A) PCA and (B) hierarchical clustering of z-scored 84-gene RT^2^ Profiler expression across untreated chondrocytes, IFP EV-treated chondrocytes and SW1353 chondrosarcoma cells (n = 3 per group); SW1353 is transcriptionally distinct while treated and untreated chondrocytes overlap. (C) Volcano plot, SW1353 vs chondrocytes and (D) IFP EVs vs untreated chondrocytes; genes with p < 0.05 coloured (red up, blue down). (E) Significantly differentially expressed genes in SW1353 vs chondrocytes. (F) Relative expression (2^-ΔCt) of VEGFA and BMP4 in untreated vs IFP EV-treated chondrocytes (box, IQR; whiskers, range; points, individual samples; two-sample t-test). (G) KEGG pathway membership among the 20 significant RT^2^ Profiler DEGs, shown as gene counts; cytokine-cytokine receptor interaction (grey) reflects the pre-curated cytokine/angiogenesis content of the RT^2^ Profiler panel. The corresponding phospho-kinase/KEGG-VEGF-pathway overlap is shown alongside the rest of the phospho-kinase array data in Figure 3D.

Against this stable chondrocyte background, IFP EVs produced a focused response. Of the 84 genes, three genes reached nominal significance: VEGFA, the central pro-angiogenic gene, was upregulated (+1.63-fold, p = 0.036); BMP4 was downregulated (–2.39-fold, p = 0.011); and FAM3B was downregulated (p = 0.032) (Figure 2D, F).

To place these transcriptional changes in pathway context, we mapped all 20 genes significantly regulated across both RT^2^ Profiler comparisons (SW1353 vs chondrocyte and IFP EVs vs untreated chondrocyte) onto the KEGG database [13] via the KEGG REST API (Figure 2G). VEGFA was the only DEG mapping to the KEGG VEGF signalling pathway (hsa04370). Cross-referencing that pathway’s gene membership against our 16 phospho-kinase array targets identified five direct overlaps: eNOS (NOS3), PLC-γ1 (PLCG1), ERK1/2 (MAPK1/MAPK3), Src and HSP27 (HSPB1) (Figure 3D), independently linking the transcriptional and phospho-protein arms of this study through a shared canonical pathway. BMP4, together with BMP2, GDF5, TGFB2 and INHBA, mapped to TGF-β signalling (hsa04350) and Hippo signalling (hsa04390), consistent with BMP4’s established anti-angiogenic role via this superfamily [14]. We also checked whether any DEGs overlapped with KEGG’s dedicated chondrosarcoma disease entry (H02902); this entry is curated around a single gene, EXT1 (heparan sulfate biosynthesis, hereditary multiple exostoses), reflecting chondrosarcoma’s known genetic aetiology rather than a broader expression signature, and none of our DEGs overlapped with it. The majority of DEGs (17/20) mapped to the generic cytokine-cytokine receptor interaction pathway (hsa04060); since the RT^2^ Profiler panel is itself a pre-curated cytokine/angiogenesis gene set, this reflects the design of the array rather than a genuine enrichment signal. KEGG pathway membership is reported here as gene-level mapping rather than a statistically corrected enrichment test, and it converges independently with the phospho-kinase data on the same VEGF-axis nodes identified in Section 3.3.

**Figure 3.**
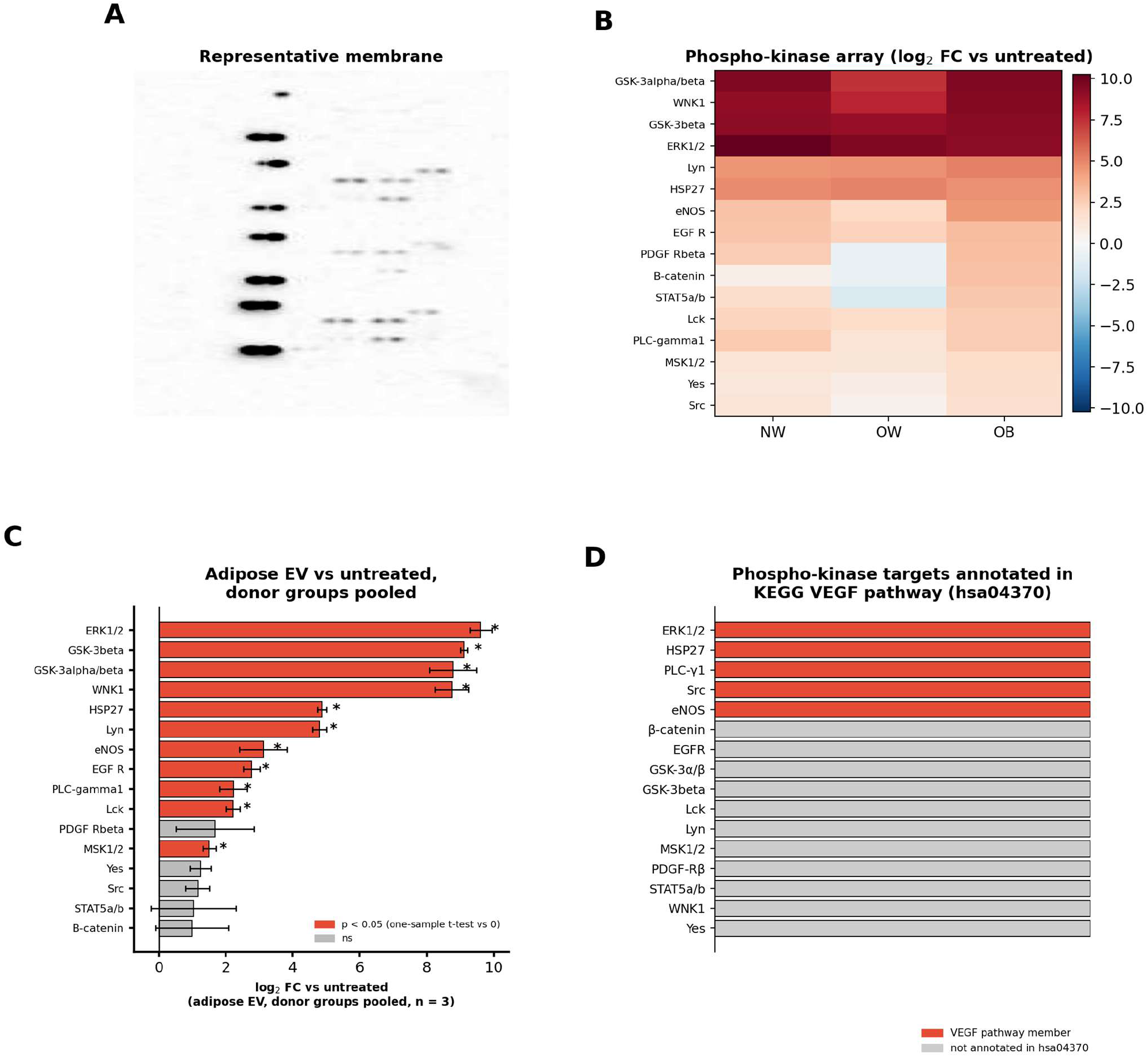
Phospho-kinase response to IFP EVs (reanalysed from Price et al. [18]), Proteome Profiler array, and its overlap with the KEGG VEGF pathway. (A) Representative Proteome Profiler membrane, cropped to the single analyte spot column; the source scan includes a duplicate reference-spot column to the right of the analyte array, which is excluded here. (B) Phospho-kinase array (log_2_ fold change vs untreated; single pooled sample per BMI group, no replicates) across normal-weight (NW), overweight (OW) and obese (OB) donor EVs, reanalysed from Price et al. [18], in which human OA primary cells were treated with the same donor IFP EV fractions used throughout this study. (C) The same 16 targets with the three BMI groups pooled as independent replicates of a single adipose EV-treated condition (n = 3), ranked by mean log_2_ fold change vs untreated; red bars reach nominal significance (one-sample t-test vs 0, p < 0.05), error bars show SEM. (D) The 16 phospho-kinase array targets grouped by whether they are annotated members of the KEGG VEGF signalling pathway (hsa04370): eNOS, PLC-γ1, ERK1/2, Src and HSP27 (red) are members; the remaining 11 targets (grey) are not.

### 3.3 Reanalysis of phospho-kinase dataset shows BMI-dependent engagement of the VEGF axis

To ask whether IFP EVs engage the VEGF-axis phospho-signalling identified in Section 3.2 more broadly, we reanalysed our previously published phospho-kinase dataset generated in human OA primary cells treated with the same normal-weight, overweight and obese donor IFP EV fractions [18] (one pooled sample per BMI group, no replicates) (Figure 3B). Cells treated with IFP EVs showed higher relative phosphorylation of the VEGF-canonical effector eNOS, alongside ERK1/2 and other targets, than untreated cells, with the highest signal seen in the obese-donor sample. Because eNOS is a direct downstream effector of VEGF signalling, this independently corroborates the KEGG VEGF-pathway overlap identified in Section 3.2 (Figure 3D).

To ask which of these targets were consistently engaged rather than driven by a single donor group, we also collapsed the three BMI groups into a single IFP EV-treated group (n = 3 donor groups, treated as independent replicates of one ‘adipose EV’ condition) and tested each target’s log_2_ fold change against zero (one-sample t-test) (Figure 3C). Eleven of sixteen targets reached nominal significance on this pooled basis, including eNOS (p = 0.048), PLC-γ1 (p = 0.033), ERK1/2 (p = 0.001) and HSP27 (p < 0.001), the same four kinases that independently map to the KEGG VEGF signalling pathway (Section 3.2, Figure 3D), reinforcing that this signal is not an artefact of the obese-donor sample alone. By contrast, β-catenin, STAT5a/b and PDGF-Rβ, which appeared prominent in the BMI-stratified view because of a large obese-donor value, did not reach nominal significance once pooled across donors (p = 0.46, 0.50 and 0.29 respectively), indicating that their signal is not consistent across donor groups. Src and Yes1 (a Src-family kinase) were similarly not significant when pooled (p = 0.084 and 0.058).

## 4. Discussion

This study shows that the IFP is a more EV-productive depot than subcutaneous fat, which, combined with its known inflammatory phenotype and direct cartilage contact, is consistent with it being a relevant adipose EV source for cartilage-lineage cells locally, including any adjacent neoplastic cartilage. Furthermore, we show that IFP EVs induce a pro-angiogenic transcriptional response in articular cartilage cells: VEGFA is upregulated, BMP4 is downregulated, and the VEGF-canonical effector eNOS, together with ERK1/2, PLC-γ1 and HSP27, is activated at the protein level, the same VEGF signalling axis reported to drive angiogenesis in chondrosarcoma via adipokine-PI3K/Akt/mTOR/HIF-α signalling [3,4]. Together these observations support a model in which joint IFP EVs can induce the angiogenic switch implicated in chondrosarcoma progression, giving a candidate cellular mechanism for adipose-driven tumour angiogenesis.

The transcriptional response is focused rather than global. PCA and clustering show that IFP EV-treated chondrocytes remain close to untreated chondrocytes and stay distinct from the chondrosarcoma line SW1353. This indicates that IFP EVs induce a targeted angiogenic signalling shift in articular cartilage-lineage cells without pushing them toward the broader dedifferentiated, malignant transcriptional state captured by SW1353, consistent with a model in which local EV signalling promotes the angiogenic switch without itself driving cellular transformation.

Such a targeted effect is more suggestive of an EV-mediated delivery of specific regulatory cargo, and points to the VEGF/eNOS axis as the primary site of impact. The concurrent reduction in expression of BMP4 is mechanistically coherent. In other tissues, BMP4 restrains angiogenesis via induction of the anti-angiogenic factor thrombospondin-1 [14], so its reduction here would be expected to remove a brake on VEGF-driven vessel growth rather than reflect an independent, unrelated change. BMP4 signalling has also been reported to support cartilage matrix synthesis, promoting type II collagen and aggrecan expression in cartilage repair models [15]. Therefore, its reduction in IFP EV-treated chondrocytes could plausibly exacerbate cartilage loss via reduced matrix production, as well as through promoting angiogenesis. Of the three genes reaching nominal significance in this comparison, FAM3B has no established role in the VEGF axis; however, it has been reported as an FGFR ligand capable of activating ERK/MAPK signalling and promoting proliferation in several cancers [16], and its broader known roles in metabolism, apoptosis and cancer warrant further investigation.

The KEGG pathway mapping adds independent, gene-level support to this chondrosarcoma link: of the 20 significantly regulated RT^2^ Profiler genes, six mapped to the generic ‘Pathways in cancer’ pathway (hsa05200) and four to PI3K-Akt signalling (hsa04151), the same signalling arm through which adiponectin and resistin drive VEGF-A-dependent angiogenesis in chondrosarcoma cells [3,4], while the phospho-kinase overlap with the KEGG VEGF pathway (eNOS, PLC-γ1, ERK1/2, Src, HSP27) sits directly downstream of this axis. This convergence across an independent transcriptional screen, a phospho-protein screen and pathway-database mapping strengthens the case that IFP-derived EVs engage the same signalling machinery implicated in chondrosarcoma angiogenesis.

The phospho-kinase screen showed the greatest eNOS, β-catenin and STAT5 signal in the obese-donor sample, raising the possibility that donor adiposity tunes the strength of this angiogenic signalling; pooling the three donor groups as replicates of one adipose EV condition supported eNOS, ERK1/2, PLC-γ1 and HSP27 as the most consistently engaged targets across donors, while β-catenin, STAT5a/b and PDGF-Rβ were not consistent once donor identity was pooled out. One plausible candidate cargo is miR-150-5p, a microRNA with an established direct suppressive action on VEGFA transcripts in other tissue contexts [17], which our group has previously shown to be transferred by adipose EVs to silence target transcripts in skeletal muscle [18]. Whether it, or another EV cargo factor, mediates the VEGFA induction reported here has not been tested and remains a hypothesis for follow-up work.

In summary, our data evidence that IFP-derived EVs provide an angiogenic signal with implications more broadly for our understanding of how non-mechanical signals from local joint adipose tissue impact on cartilage pathology [19], and, more specifically, how a pathological VEGF pro-angiogenic signal could mediate chondrosarcoma.

## 5. Limitations

VEGFA and BMP4 were nominated from an 84-gene screen using nominal, uncorrected p-values. The phospho-kinase array was likewise designed as a pathway-nomination tool rather than a fully replicated signalling study. Independent qPCR confirmation in an expanded donor cohort, and quantitative repeats of the phospho-kinase readout, are the natural next steps but beyond the scope of this paper. The IFP-versus-subcutaneous EV comparison used independent rather than donor-paired samples, and the EV cargo responsible for VEGFA induction remains to be identified. The chondrosarcoma-relevant conclusions drawn here rest on comparisons with a single immortalised chondrosarcoma cell line (SW1353) rather than primary human chondrosarcoma tissue, and the proposed fat pad EV-chondrosarcoma link has not been tested directly in a chondrosarcoma model.

## Data availability

Data are available from the corresponding author upon reasonable request.

## Acknowledgements and funding

This study was supported by the Dubrowsky Legacy Foundation, MyAge, the UKRI Medical Research Council (grant reference MR/W026961/1), and Arthritis UK (grant reference 21812). The authors would like to thank the Dubrowsky Legacy Foundation for funding and support for J.M.J.P. The authors would like to thank the Oxford Brookes Centre for Bioimaging where transmission electron microscopy was performed. This study was delivered through the National Institute for Health and Care Research (NIHR) Birmingham Biomedical Research Centre (BRC), and supported by NIHR award NIHR-INF-5081. The views expressed are those of the author(s) and not necessarily those of the BRC, the NIHR, the Department of Health and Social Care, or any of our other funders.

## Conflict of interest

The authors declare no conflicts of interest.

## Notes

### Competing Interest Statement

The authors have declared no competing interest.

