## Supplementary Materials for "Infrapatellar Fat Pad Extracellular Vesicles Induce a Pro-Angiogenic VEGFA^high^/BMP4^low^ Switch in Articular Chondrocytes: Implications for Chondrosarcoma"

*Table S1. MIFlowCyt-EV reporting framework for the nanoscale flow cytometry EV concentration assay (CytoFLEX nano, Beckman Coulter; Section 3.1, Figure 1A). Items are reported as described in Methods 2.2 and Results 3.1.*

| Component | Status | Details |
|---|---|---|
| 1.1 Preanalytical variables (MISEV) | Reported | Tissue mincing, 24 h DMEM conditioning (37°C, 21% O2, 5% CO2), clarification (2,000×g ×2) and 0.22 µm filtration described in Methods 2.2; study follows MISEV2023 guidance [11]. |
| 1.2 Experimental design (MIFlowCyt) | Reported | Aim/hypothesis and comparison defined in Section 3.1: fat pad vs subcutaneous ACM EV concentration, n = 8 donors per depot, one-sided Student's t-test (fat pad predicted a priori higher, given its known inflammatory phenotype [7,8]). |
| 2.1 Sample staining details | Not applicable | Label-free, scatter-based detection; no fluorescent antibody or dye staining was used for EV enumeration. |
| 2.2 Sample washing details | Not applicable | No post-collection wash step; ACM was clarified and 0.22 µm-filtered prior to analysis (Methods 2.2). |
| 2.3 Sample dilution details | Reported | EV samples (fat pad and subcutaneous ACM) were diluted 1:400–1:800 to remain within the linear detection range of the instrument, consistent with the dilution-linearity validation ($R^2$ = 0.997) (Methods 2.2). |
| 3.1 Buffer-only controls | Reported | A buffer-only control of 0.22 µm-filtered PBS was acquired alongside samples at matched CytoFLEX nano acquisition settings. |
| 3.2 Buffer with reagent controls | Not applicable | No reagents or antibodies were added to samples. |
| 3.3 Unstained controls | Not applicable | No staining was performed. |
| 3.4 Isotype controls | Not applicable | No antibody staining was performed. |
| 3.5 Single-stained controls | Not applicable | No multiplexed fluorescent staining was performed. |
| 3.6 Procedural controls | Reported | Procedural controls (processed identically to EV samples but without EV-containing material) were run alongside samples and returned a blank (background-level) signal. |
| 3.7 Serial dilutions | Reported | Dilution-linearity validation performed ($R^2$ = 0.997), supporting single-particle (non-swarm) detection; swarm-detection controls also applied (Methods 2.2). |
| 3.8 Detergent-treated controls | Not performed | Detergent (e.g. Triton X-100) lysis controls were not performed for this assay. |
| 4.1 Trigger channel(s)/threshold(s) | Reported | Triggered on violet side scatter (VSSC1); threshold = 250 (arbitrary units). |
| 4.2 Flow rate/volumetric quantification | Reported | Flow rate was set at 1 µL/min on the CytoFLEX nano. |
| 4.3 Fluorescence calibration | Not applicable | No fluorescent labelling was used; fluorosphere calibration beads were used for light-scatter/size referencing (Methods 2.2), not for fluorescence intensity. |
| 4.4 Scatter calibration | Not performed | No scatter calibration (e.g. Mie-theory-based scatter-to-$nm^2$ conversion) was performed for this assay; the VSSC1 trigger threshold (Component 4.1) was used in arbitrary units. |

| Component | Status | Details |
|---|---|---|
| 5.1 EV diameter/surface area/volume | Not applicable to this assay | Particle sizing was performed by nanoparticle tracking analysis (NTA; NanoSight Pro), not by nanoscale flow cytometry, in this study (Figure 1B, C, E). |
| 5.2 EV refractive index approximation | Not applicable | Not determined by flow cytometry in this study. |
| 5.3 EV epitope number approximation | Not applicable | No immunophenotyping was performed; this assay measured concentration only. |
| 6.1 Completion of MIFlowCyt checklist | Reported (this table) | This MIFlowCyt-EV-based summary constitutes the completed checklist for this assay; a separate instrument-operator MIFlowCyt v1.0 form was not appended. |
| 6.2 Calibrated channel detection range | Not reported | Not determined; no scatter calibration was performed to convert the detection range into standardised units (Component 4.4). |
| 6.3 EV number/concentration | Reported | Fat pad ACM: mean $1.42\times10^{11}$ particles/mL; subcutaneous ACM: mean $5.66\times10^{10}$ particles/mL ($n = 8$ per depot; one-sided t-test $p = 0.027$) (Section 3.1, Figure 1A). |
| 6.4 EV brightness | Not applicable | No fluorescence was measured; concentration-only assay. |
| 7.1 Sharing of data to a public repository | Reported (partial) | Source data are available from the corresponding author on request (Data availability statement); not deposited in a public repository (e.g. FlowRepository). |
